# Corporately Owned Plants need Space: An Analysis on Plant Biodiversity in Toronto’s Privately-owned Publicly Accessible Spaces

**DOI:** 10.1101/2023.01.29.526117

**Authors:** Matthew R Ho

## Abstract

Urban plant biodiversity is a growing ecological concern for city planners and ecologists. While parks are serviced by the public sector, and yards are pruned by the private citizen, a growing proportion of urban green space is managed by corporations. Despite biodiversity targets set by city councils and public committees, actual plant surveys have not been performed. Employing coordinate data of Privately-Owned Publicly-accessible Spaces (POPS) from the City of Toronto, we sampled plant species richness in nine corporately-managed green spaces. Using linear mixed-effect models, we compared richness with various green space characteristics and found that site area is an important predictor. Our results concur with prior studies showing that habitat area may cause significant impacts to urbanized plant biodiversity.

## Introduction

To provide residents with much-needed leisure space, the development of Privately-Owned Publicly-accessible Spaces (POPS) is an increasingly common request from urban planning and development boards. In the City of Toronto, building developers negotiate for building code exemptions (such as increased floor area, taller maximum height, reduced building setbacks, etc.) in exchange for private funding towards city parks, recreation, or the creation of POPS on their development footprint (Schmidt et al. 2011, Moore 2013). Established in 2013, Toronto POPS guidelines can be functionally classified into six categories: courtyards, plazas, gardens, forecourts, sidewalk landscaping, and pedestrian walkways. To reach the city’s sustainability goals, recommended POPS landscaping is required meet tier 1 of the Toronto Green Standard (TGS) (City of Toronto 2012). The TGS was introduced in 2006 to address air quality, energy usage, storm-water runoff, and greenhouse gas emissions. Starting as a voluntary program, version 4 of the TGS is now mandatory for all new building developments, with different guidelines for low-rise and high-rose buildings. The mandatory tier 1 – the lowest tier – obliges POPS to plant at least 50 percent of native vegetation (Nievas 2019). Albeit, various culturally important non-native and invasive species are on the official list of species recommendations. The importance of species compositions cannot be understated. Prior research has demonstrated a shift in insect species at different proportions of native/non-native vegetation (Beninde et al. 2015).

Since the formation of official guidelines, Toronto POPS built before 2013 are slowly being indexed (City of Toronto 2014). Few records of landscaping requirements can be found on these properties. As such, a systematic sample of plant species compositions is an important first step to elucidate the condition of these green spaces (Silva and Stefani 2018). In our study, we survey POPS sites from properties of varying location, age, and height. Then, we examine if site characteristics have any effect on the proportion of site-level native vegetation. Collectively, this project serves as a starting point for more POPS analyses around the world. Moreover, this project provides insight for city planners as to which policies closely affects species composition. All things considered, any analysis performed adds valuable discussion on the eco-dynamics of this ever-urbanizing world.

### Hypothesis

To investigate the relationship between plant species composition and POPS characteristics, mixed-effect models are used. While there are many characteristics to explore, selected elements include site age, site area, building height, neighbourhood income, and surrounding population density. Our hypothesis is as such:

Alternate hypotheses (HA): POPS characteristics have a significant effect on the percentage of native species composition.
Null hypothesis (H0): POPS characteristics do not have a significant effect on the percentage of native species composition.

These elements were selected due to their likelihood of producing an effect. First, site age may indicate a dichotomy between sites established before, and after 2013. Next, increased site area may encourage landscapers be creative in plantings. Third, building height negotiations may incentivize developers to closely follow city guidelines. Fourth, neighbourhood income may affect the developer’s landscaping choices. Finally, surrounding population density may influence building development configurations.

## Methods

### Site Selection

The scattered distribution of Toronto POPS requires random selection for an accurate sampling. Since 2013, a list of all new, inducted, and future POPS is displayed as an interactive map on the city website. We extracted this dataset from the City of Toronto’s Open Data portal and grouped each site by neighbourhood. Since POPS vary in category, we filtered for sites with “parkette”, “garden”, “open space”, or “landscaping” in their site description. This removed sites with little vegetation such as walkways or forecourts. Using the function “sample_frac” in R package dplyr, we randomly selected 40 percent of all sites to sample (v 1.0.10, Wickham 2022). This was done in R version 4.2.1. To visualize the sites for easy sampling, we mapped the subset using leaflet (v 2.1.1, Cheng 2022).

Since POPS generally occur in small patches, the tried-and-true transect method is impossible. Moreover, since preliminary field surveys indicate many POPS exhibit a low species variety, sampling species richness for the entire site is feasible. As such, field sampling consists of walking the entire site and recording all unique species. To aid in species recognition, computer algorithms, such as Google Lens and Seek by iNaturalist, were used (Anderson 2018).

Site characteristics were extracted from the 2016 Canadian Census dataset and the emporis.com database. Specially, we extracted neighbourhood income (“v_CA16_2201”) and surrounding population density (“v_CA16_406”) using package cancensus (v 0.5.4, Shkolnik 2022). Building height was extracted from the website emporis.com. This is a real-estate development website with a large back-end database normally inaccessible by regular web-users. To access the data, any search engine’s cached webpage data can be used. We searched the address with the search engine’s site search condition set to site:emporis.com. Site age was also estimated based on the building’s construction year or the building’s most recent renovation year. Finally, POPS site area was estimated by drawing polygons in satellite aerial view from a GIS software. The site area estimate was based on the soil area for each POPS. Impervious surfaces were not included.

To add the extracted POPS characteristics to our collected data, we used the function “geocode” in package ggmap to generate coordinates of each site (v 3.0.1, Kahle 2022). Next, we use the function “st_join” in package spatial points to locate and append from which census neighbourhood each site is located (v 1.5-1, Pebesma 2022). Finally, we used “left_join” in dyplr to merge all POPS characteristics to our collect data. After pivoting from wide to long, the data can be analyzed.

### Statistical Analyses

To analyze if site characteristics have any effect on species composition, we employed linear mixed-effect models with the percentage of native species sampled as the dependent variable. Site area, building height, POPS age, income, and density were included as fixed effects. All models were performed using the “lmer” function in package lme4 (v 1.1-31, Bolker 2022). To account for the possible effect that commercial or residential function has the model, we included the building type as a random variable.

Our linear effect models were performed in a stepwise manner. We performed individual models with area, height, age, income, and density as separate functions. We also performed a combined function with all fixed effects. This is our based model To refine our base model, we isolated the three fixed-effect variables with the lowest p-values to run another model. These variables were site area, building height, and POPS age. To further explain the area model, we used the function “dredge” from package MuMIn (v1.47.1, Bartoń 2022) to perform a dredge of this model. The dredge function performs automated model selection to see which predictors, or combination of predictors, is the best explanation for the data. Finally, we used the function “model.avg” from package MuMIn to obtain estimated coefficients for the fixed effects based of the selected models.

## Results

### Detailed Dataset

The estimates of coefficients described by the fixed effects in the linear-mixed models “Estimate” as well as the corresponding z-stats “z” and p-val “p” were obtained from the results of the averaged linear mixed models for each of the three morphological traits (see table 1 for base model). Our initial base models produced no significant effects; however, site area contained the lowest p-value. Our refined model produced a statistically significant result for site area (Estimate = 0.314, *z* = 0.1305, *p* = 0.0457) (see table 2 for area model results). After performing a dredge, surprisingly, most significant with lowest Akaike information criterion (AICc) predictor was POPS age (see table 3).

To test for collinearity with site area, we performed a fixed-effect model with site area as our dependent variable. We used all the other variables from our base model, as predictors. These predictors were set to have a crossed interaction term with all other predictors. No significant results were produced (*p* > 0.05) indicating no correlation between site area and the other POPS characteristics (see table 4).

### Statistical Limitations

It is important to recognize that linear models on small dataset of real-world data can be extremely ineffective. Our refined native ~ area model demonstrates this concept well; notably, if building height and POPS age were removed, the model would return not statistically significant. Although our test for correlation to area was nonsignificant, it is still possible that height and age plays some part in POPS area. Further analyses can incorporate a Sobel test to understand the mechanism that underlies the relationship between these three variables (Baron and Kenny 1986).

## Discussion

Our results seem to loosely support our hypothesis. While there is statistically significant support for increased native proportion with increased area, some interplay between our predictors cast doubt on this result. In fact, many of the chosen predictors did not produce a significant effect on the percentage of native vegetation. This indicates that the effect of each POPS characteristic can be distinct to specific locations and environmental conditions, and are not generalized across the study area. Additionally, since our sites are limited in spatial scope, it is difficult to draw concrete trends in the data. Below, we present a few theories on why the selected characteristics may be erroneous.

### Effect of Income and Density

Although income and density were extracted from the 2016 Canadian census at a very fine detail (by the city block), POPS plants are managed by corporations that do not integrate into the surrounding socioeconomic landscape (Rigolon and Németh 2018). This is to say that corporately managed green-space may be highly homogenous over the economic spectrum. Further studies on POPS homogeneity in general should be pursued. This may include studies on the correlation between architectural design and hard landscaping to the vegetation composition found at each site. There is evidence to suggest that urban design aspect plays a role in the distribution of urban greenery; specifically, lower income neighbourhoods having decreased plant biomass (Wolch et al. 2014). A meta-analysis on public greenery and private greenery across different economic spectra may elucidate exciting information.

### Effect of Building Height and Age

While our estimates indicate some correlation between native species composition and building characteristics, our method of determining these attributes may be flawed. The ideal method is to access the city’s building records, requiring a fee for each record accessed. City records will need to be combed for the year of which that POPS was created. Private organizations may also have built their respective POPS without registering or notifying city hall. Moreover, the expectation from municipal planning board negotiations may also be highly variable from site to site; for example, where one developer might offer large green spaces, another may offer affordable housing units. Finally, the effect of POPS age may be heavily skewed if the corporation changes landscaping strategies, or if the landscaping company replaces plant species over time. Without accessing company internal documents, it may be impossible to know the true effect of POPS age. Future studies should exclude these variables, and instead investigate the effect of the current landscaping company on native vegetation (Dromgold et al. 2020).

### Effect of Site Area

Regarding the percentage of native vegetation per site, site area produces the greatest effect. Future studies require a finer area measurement using ground-level tools, such as a measuring wheel. Coarse-level GIS data may be highly inaccurate when measuring the smallest POPS, and thus grossly overestimate some regions. However, there is significant evidence that the area of publicly-managed urban green-spaces plays a role in biodiversity (Beninde et al. 2015). According to a meta-analysis on public green-spaces, site area and site water cover were the largest abiotic factor in determining biodiversity. Further research on privately-owned site area may explain biotic trends; however, our data moderately agrees with the consensus.

## Conclusion

Our original hypothesis highlights that POPS characteristics can affect native plant species composition. The results indicate that site area has a significant impact, with building height and POPS age playing some role. Our analyses lead us to reject the null hypothesis of POPS characteristics not significantly affecting native species compositions. Though our data collection can be improved, our models concur with prior research on urban biodiversity illustrating that biota need space to thrive.

## Appendix 1: Tables

**Table 1.** These are the base model results with all predictors included. The model formula is as follows: (perc.native ~ site.area + build.height.ft + POPS.age + income + density + (1 | build.type)). Building type is included as a random variable. There is no statistically significant predictor in our base model.

|  | Estimate | z-val | p-val |
| --- | --- | --- | --- |
| (Intercept) | 3.283e-01 | 1.822e-01 | 0.803 |
| site.area | 3.623e-05 | 2.630e-05 | 0.277 |
| build.height.ft | 2.238e-05 | 1.827e-04 | 0.913 |
| POPS.age | -3.078e-03 | 5.722e-03 | 0.744 |
| income | -7.020e-05 | 8.400e-05 | 0.465 |
| density | 3.225e-06 | 3.960e-06 | 0.976 |

**Table 2.** These are the refined model results, with the model formula as follows: (perc.native ~ site.area + build.height.ft + POPS.age + (1 | build.type). Building type is again included as a random variable. In the refined model, there is slight statistical significance for the area predictor (p < 0.05). This result concurs with prior research on urban biodiversity.

|  | Estimate | z-val | p-val |
| --- | --- | --- | --- |
| (Intercept) | 3.146e-01 | 1.305e-01 | 0.0608 |
| site.area | 4.869e-05 | 1.841e-05 | 0.0457* |
| build.height.ft | 7.115e-05 | 1.266e-04 | 0.5984 |
| POPS.age | -5.983e-03 | 3.952e-03 | 0.1904 |

**Table 3.** Model selection table using the function “dredge” on the refined area model. Of note in this table is the Akaike information criterion (AICc) values indicating which predictor performs the best. The lowest AICc value is shown to be none, with POPS age following. This indicates some interplay between POPS age, build height, and site area.

| Model selection table |  |  |  |  |  |  |  |  |  |
| --- | --- | --- | --- | --- | --- | --- | --- | --- | --- |
|  | (Int) | bld.hgh.ft | POP.age | sit.are | df | logLik | AICc | delta | weight |
| 1 | 0.2217 |  |  |  | 3 | 2.910 | 5.0 | 0.00 | 1 |
| 3 | 0.3053 |  | -0.002430 |  | 4 | -1.421 | 20.8 | 15.86 | 0 |
| 2 | 0.2434 | -4.739e-05 |  |  | 4 | -4.925 | 27.8 | 22.87 | 0 |
| 5 | 0.1581 |  |  | 3.843e-05 | 4 | -5.019 | 28.0 | 23.06 | 0 |
| 7 | 0.3263 |  | -0.005228 | 4.561e-05 | 5 | -8.687 | 47.4 | 42.39 | 0 |
| 4 | 0.3106 | -2.854e-05 | -0.002203 |  | 5 | -9.200 | 48.4 | 43.42 | 0 |
| 6 | 0.1551 | 5.982e-06 |  | 3.860e-05 | 5 | -13.076 | 56.2 | 51.17 | 0 |
| 8 | 0.3146 | 7.115e-05 | -0.005983 | 4.869e-05 | 6 | -16.606 | 87.2 | 82.23 | 0 |

**Table 4.** These are the results of the linear model with site area as the dependent variable. This model was created to test if any of the general predictors had any correlation with site area. No correlation was found indicating little collinearity. A principal component analysis may be useful in this case.

|  |  |  |  |  |  |  |  |  |
| --- | --- | --- | --- | --- | --- | --- | --- | --- |
| 1 | 2 | 3 | 4 | 5 | 6 | 7 | 8 | 9 |
| -2.754e-13 | 2.812e-12 | -2.406e-13 | -1.829e+00 | 6.184e+01 | -5.063e+02 | 4.677e+02 | -2.141e+01 | 7.466e-13 |

Coefficients: (8 not defined because of singularities)
|  | Estimate | Std. Error | t value | Pr(>\|t\|) |
| --- | --- | --- | --- | --- |
| (Intercept) | 1.319e+05 | 2.417e+04 | 5.458 | 0.1154 |
| POPS.age | -2.105e+03 | 4.585e+02 | -4.591 | 0.1365 |
| build.height.ft | -2.415e+02 | 3.964e+01 | -6.092 | 0.1036 |
| income.CA16.2201 | -1.183e+02 | 1.992e+01 | -5.938 | 0.1062 |
| density.CA16.406 | 8.125e-01 | 1.213e-01 | 6.696 | 0.0944 |
| POPS.age:build.height.ft | 6.110e-02 | 1.314e-01 | 0.465 | 0.7229 |
| POPS.age:income.CA16.2201 | 1.824e+00 | 3.354e-01 | 5.440 | 0.1157 |
| build.height.ft:income.CA16.2201 | 2.155e-01 | 3.360e-02 | 6.414 | 0.0985 |
| POPS.age:density.CA16.406 | NA | NA | NA | NA |
| build.height.ft:density.CA16.406 | NA | NA | NA | NA |
| income.CA16.2201:density.CA16.406 | NA | NA | NA | NA |
| POPS.age:build.height.ft:income.CA16.2201 | NA | NA | NA | NA |
| POPS.age:build.height.ft:density.CA16.406 | NA | NA | NA | NA |
| POPS.age:income.CA16.2201:density.CA16.406 | NA | NA | NA | NA |
| build.height.ft:income.CA16.2201:density.CA16.406 | NA | NA | NA | NA |
| POPS.age:build.height.ft:income.CA16.2201:density.CA16.406 | NA | NA | NA | NA |

## References

Anderson, S. W. 2018. iNaturalist: Understanding Biodiversity Through a Digital Medium.

Baron, R. M., and D. A. Kenny. 1986. The Moderator-Mediator Variable Distinction in Social Psychological Research. Conceptual, Strategic, and Statistical Considerations. Journal of Personality and Social Psychology 51:1173–1182.

Beninde, J., M. Veith, and A. Hochkirch. 2015. Biodiversity in cities needs space: a metaanalysis of factors determining intra-urban biodiversity variation. Ecology Letters 18:581–592.

City of Toronto. 2012. DROUGHT TOLERANT LANDSCAPING A Resource for Development.

City of Toronto. 2014. Creative Place Making to Enhance Urban Life PRIVATELY-OWNED PUBLICLY ACCESSIBLE SPACES.

Dromgold, J. R., C. G. Threlfall, B. A. Norton, and N. S. G. Williams. 2020. Green roof and ground-level invertebrate communities are similar and are driven by building height and landscape context. Journal of Urban Ecology 6.

Moore, A. 2013. Trading Density for Benefits: Section 37 Agreements in Toronto. Institute on Municipal Governance and Finance, Munk School of Global Affairs, University of Toronto 14.

Nievas, M. 2019. Planning Healthy Cities: Privately-owned publicly-accessible spaces in Toronto.

Rigolon, A., and J. Németh. 2018. Privately owned parks in New Urbanist communities: A study of environmental privilege, equity, and inclusion. Journal of Urban Affairs 40:543–559.

Schmidt, S., J. Nemeth, and E. Botsford. 2011. The evolution of privately owned public spaces in New York City 16:270–284.

Silva, F., and M. Stefani. 2018. Biodiversity urban: how the city can do its management?

Wolch, J. R., J. Byrne, and J. P. Newell. 2014. Urban green space, public health, and environmental justice: The challenge of making cities ‘just green enough.’ Landscape and Urban Planning 125:234–244.

